# Genetic analysis of *de novo* variants reveals sex differences in complex and isolated congenital diaphragmatic hernia and indicates *MYRF* as a candidate gene

**DOI:** 10.1101/206037

**Authors:** Hongjian Qi, Lan Yu, Xueya Zhou, Alexander Kitaygorodsky, Julia Wynn, Na Zhu, Gudrun Aspelund, Foong Yen Lim, Timothy Crombleholme, Robert Cusick, Kenneth Azarow, Melissa Ellen Danko, Dai Chung, Brad W. Warner, George B. Mychaliska, Douglas Potoka, Amy J. Wagner, Mahmoud ElFiky, Deborah A. Nickerson, Michael J. Bamshad, Jay M. Wilson, Frances A. High, Mauro Longoni, Patricia Donahoe, Wendy K. Chung, Yufeng Shen

**Affiliations:** Department of Systems Biology, Columbia University, New York, NY, USA; Department of Applied Mathematics and Applied Physics, Columbia University, New York, NY, USA; Department of Pediatrics, Columbia University, New York, NY, USA; Department of Biomedical Informatics, Columbia University, New York, NY 10032; Department of Surgery, Columbia University Medical Center, New York, NY, USA; Cincinnati Children's Hospital, Cincinnati, OH, USA; Children's Hospital & Medical Center of Omaha, University of Nebraska College of Medicine, Omaha, NE, USA; Department of Surgery, Oregon Health&Science University, Portland, OR, USA; Monroe Carell Jr. Children's Hospital, Vanderbilt University Medical Center, Nashville, TN, USA; Washington University, St. Louis Children's Hospital, St. Louis, MO, USA; University of Michigan, CS Mott Children's Hospital, Ann Arbor, MI, USA; Children's Hospital of Pittsburgh, Pittsburgh, PA, USA; Medical College of Wisconsin, Milwaukee, WI, USA; Department of Pediatric Surgery, Faculty of Medicine, Cairo University, Cairo, Egypt; Center for Mendelian Genomics, University of Washington, Seattle, Washington, USA; Department of Surgery, Boston Children’s Hospital, Boston, MA, USA; Department of Surgery, Harvard Medical School, MA, USA; Pediatric Surgical Research Laboratories, Department of Surgery, Massachusetts General Hospital, Boston, MA, USA; Department of Medicine, Columbia University, New York, NY, USA; Herbert Irving Comprehensive Cancer Center, Columbia University, New York, NY, USA; JP Sulzberger Columbia Genome Center, Columbia University, New York, NY, USA

## Abstract

Congenital diaphragmatic hernia (CDH) is one of the most common and lethal birth defects. Previous studies using exome sequencing support a significant contribution of coding *de novo* variants in complex CDH cases with additional anomalies and likely gene-disrupting (LGD) variants in isolated CDH cases. To further investigate the genetic architecture of CDH, we performed exome or genome sequencing in 283 proband-parent trios. Combined with data from previous studies, we analyzed a total of 357 trios, including 148 complex and 209 isolated cases. Complex and isolated cases both have a significant burden of deleterious *de novo* coding variants (1.7~fold, p= 1.2×10^−5^ for complex, 1.5~fold, p= 9.0×10^−5^ for isolated). Strikingly, in isolated CDH, almost all of the burden is carried by female cases (2.1~fold, p=0.004 for likely gene disrupting and 1.8~fold, p= 0.0008 for damaging missense variants); whereas in complex CDH, the burden is similar in females and males. Additionally, *de novo* LGD variants in complex cases are mostly enriched in genes highly expressed in developing diaphragm, but distributed in genes with a broad range of expression levels in isolated cases. Finally, we identified a new candidate risk gene *MYRF* (4 *de novo* variants, p-value=2×10^−10^), a transcription factor intolerant of mutations. Patients with *MYRF* mutations have additional anomalies including congenital heart disease and genitourinary defects, likely representing a novel syndrome.

## 1 Introduction

Congenital diaphragmatic hernia (CDH) is an anatomical defect of the diaphragm that leads to the protrusion of abdominal viscera into the thoracic cavity, compressing the lungs in utero and resulting in lung hypoplasia. CDH affects approximately 1 in 3000 live births and is often lethal^1; 2^. It can be isolated (50–60%) or associated with other birth defects and neurodevelopmental disorders ^3; 4^. Among these, cardiovascular malformations are the most common (~35% of CDH patients)^5^. Lung hypoplasia and the associated pulmonary hypertension are the main cause of the mortality and morbidity of CDH. Despite greatly improved survival rate with neonatal and surgical interventions, the overall mortality remains at ~30% ^6–8^.

The diaphragm develops between the fourth and tenth weeks of human gestation and in mice between embryonic day (E) 10.5 and E15.5^9^. Both environmental and genetic factors have been implicated. The mesenchymal-derived pleuroperitoneal folds (PPFs) play a key role in diaphragm development, and mutations in PPF-derived muscle connective tissue fibroblasts can result in CDH ^10^. Most genes implicated in CDH have been identified through recurrent chromosomal anomalies and mutant mice^11–17^. The etiology is unclear for most CDH patients. The historical low reproductive fitness of CDH has limited the number of familial cases for genetic analysis. We and others have reported an enrichment of *de novo* deleterious genetic events in sporadic CDH patients^18–20^, especially LGD (likely gene disrupting) variants in complex cases.

To identify novel risk genes and compare the genetic architecture of complex and isolated cases, we performed whole exome sequencing (WES) in 79 proband-parent trios and whole genome sequencing (WGS) in 192 trios. Combined with previously published cases^18; 19^, we analyzed a total of 357 trios (Supplementary Table 1), including 148 complex and 209 isolated cases. We observed that there are different contribution from *de novo* variants in female and male CDH cases, and genes implicated by LGD variants in complex and isolated CDH cases have distinct expression patterns in early diaphragm development. Finally, we identified *MYRF* as a new candidate risk gene with *de novo* variants in four complex CDH patients.

## Material and Methods

### Patients

A total of 357 CDH patients and their unaffected parents were recruited for analysis in this study, including 74 trios from Boston Children’s Hospital (BCH) and Massachusetts General Hospital (MGH)^18^ (Boston Cohort) and 39 trios from a previous study ^19^ (Supplementary Table 1). Two hundred and eighty-three trios were recruited as part of the DHREAMS (Diaphragmatic Hernia Research & Exploration; Advancing Molecular Science) study (http://www.cdhgenetics.com/)^20^. Neonates, children and fetal cases with a diagnosis of diaphragm defects were eligible for DHREAMS. Clinical data were abstracted from the medical chart by study personnel at each of 16 clinical sites. Data on prenatal history, neonatal outcome, and longitudinal follow-up data including Bayley III and Vineland II developmental assessments and a parent interview about the patient's health since discharge at 2 years of age and/or 5 years of age were gathered in our birth cohort. A complete family history of diaphragm defects and major malformations was collected on all patients by a single genetic counsellor, and no patients had a family history of CDH.

Patients without additional birth defects or neurodevelopmental disorder (NDD) at last contact were classified as isolated, and patients with the additional birth defects or NDD were classified as non-isolated (Details previously published^19; 20^). The diaphragm lesion was classified as left, right, bilateral or central. Pulmonary hypoplasia, cardiac displacement and intestinal herniation were considered to be part of the diaphragm defect sequence and were not considered to be an additional malformation. Subjects from BCH and MGH were described previously^18^. A blood, saliva, and/or skin/diaphragm tissue sample was collected from the affected patient and both parents. All participants provided informed consent/assent for participation in this study, which was approved by the institutional review boards of each participate study site.

### Whole exome and whole genome sequencing

We included previously two sets of WES data for analysis^18; 19^. We performed whole exome sequencing (WES) at the University of Washington in 79 additional trios using genomic DNA largely from whole blood (73 trios, 93.4%), with a minority from saliva or tissues. DNA was processed with the Nimblegen SeqCap EZ Exome V2 exome capture reagent (Roche) and TruSeq DNA Sample Prep Kits (Illumina). Samples were multiplexed and sequenced with paired-end 75bp reads on Illumina HiSeq 2500 platform according to the manufacturer’s instructions (Illumina, Inc, San Diego, California, USA).

We sequenced another 192 trios at Baylor College of Medicine using whole genome sequencing (WGS) as part of NIH Gabriella Miller Kids First Pediatric Research Program. Among these, 27 trios that had no damaging *de novo* variants in previously published WES data were selected as “WES-negative” cases for WGS^19^. Genomic libraries were prepared by the Illumina TruSeq DNA PCR-Free Library Prep Kit. DNA was sheared into 350-bp average length using sonication on a Covaris LE220 instrument. The fragmented DNA was end-repaired, A-tailed and indexed using TruSeq Illumina adapters with overhang-T added to the DNA. The libraries were validated on a Bioanalyzer DNA High Sensitivity chip by size and quality, then pooled in equal quantities and sequenced as paired-end reads of 150-bp lengths on an Illumina HiSeq X platform.

### Alignment and quality controls

Mapping, alignment, and variant calling were done according to the Broad Institute’s best practices using Burrows-Wheeler Aligner (bwa-mem, version 0.7.10)^21^ and Genome Analysis Toolkit (GATK;version3.3) (https://software.broadinstitute.org/gatk/best-practices/). Briefly, we mapped WES or reads to the reference genome (build GRCh37) using BWA-mem ^22^, mark PCR duplicates using Picard (v1.67), performed local realignment and quality recalibration using GATK ^23^. We jointly called variants in all WES samples using the GATK HaplotypeCaller. The output file was generated in the universal variant call format (VCF). We used the same procedure to analyze WGS samples.

Among new samples sequenced by WES, the mean depth of coverage is 59± 21 with 93±2.5% bases read with at least 15x in target regions. Among new samples sequenced by WGS, the mean depth of coverage is 39±2, with 99±0.25% bases read at least 15x (Supplementary Fig. 1).

We performed principal component analysis of common variants (allele frequency >5%) using Eigenstrat ^24^ to determine the population structure and ancestry of both cases and controls, with HapMap 3 sample collection data ^25^ as a reference.

### Detection of *de novo* SNVs and indels

We used Plink^26^ (http://pngu.mgh.harvard.edu/purcell/plink/) to estimate Identity by Descent (IBD)^27^ to confirm the relatedness among familial trios. All trios were matched to parents-offspring with relatedness.

A variant that presents as a heterozygous genotype in the offspring and homozygous reference genotypes in both parents was considered to be a potential *de novo* variant. We used an established stringent filtering method to identify *de novo* variants as described previously ^19; 28; 29^. Briefly, we required the candidate variants have depth (minimum 5 alternate allele reads), alternate allele fraction (minimum 20%), Fisher Strand (FS) (maximum 25), Quality by depth (QD) (minimum 2), Phread-scaled genotype likelihood (PL) (minimum 60), population allele frequency(maximum 0.1% in ExAC), and parental read characteristics (minimum depth of 10 reference reads; alternate allele fraction less than 5%, minimum GQ of 30). Additionally, variants located in segmental duplication regions (maximum score 0.98) were excluded. All candidate *de novo* variants were manually inspected in the Integrated Genomics Viewer (IGV, http://software.broadinstitute.org/software/igv/). In addition, we validated all the *de novo* likely gene disrupting (LGD) (including frameshift, nonsense and splicing site) variants by dideoxynucleotide sequencing. Of 40 case variants that were submitted for validation by Sanger sequencing, all 40 were confirmed (precision =100%).

Among the 27 “WES-negative” cases, there were 12 *de novo* variants identified by WGS that were not detected by WES ^19^.

### Annotation of variants

We used ANNOVAR^30^ to annotate variants and aggregate allele frequency and *in silico* functional predictions, then used average allele frequency in Exome Aggregation Consortium (ExAC) data to define rare variants (frequency < 1e-4). Rare *de novo* variants were classified as silent, missense, and likely-gene-disrupting (“LGD”, which includes stopgain, stoploss, canonical splicing site, or frameshift variants). In-frame insertions or deletions were not considered in the genetic analysis. We defined deleterious missense variants (“D-mis”) by CADD^31^ phred-scale score ≥25.

### Statistical analysis

We performed statistical analyses using R package from the Comprehensive R Archive Network, and the denovolyzerR ^32^ package.

### Global or gene set burden between case and mutation background rate

We calibrated the expected number of *de novo* variants in patients in each variant class in each gene based on the 3-nucleotide context-specific mutation rate estimated by Samocha et al.^29; 32^.

We used Poisson test to assess the significance of excess of observed *de novo* variants over expectation which was defined as enrichment rate (r). The positive predictive value (PPV) for *de novo* variants in each class was calculated as (*r−*1)/*r*. The Estimated number of true risk variants in each class is the number of observed variants (*m*) in cases multiplied by PPV: m * (r−1)/ r. The most severe predicted functional effect variants (LGD and D-mis) were used in further burden analyses based on the different phenotype, gender, gene set, and expression data.

### Percent of CDH attributable to *de novo* variants

We calculated the percent of CDH patients with pathogenic variants in isolated and complex CDH groups, in male and female case groups, respectively. The fraction of individuals carrying at least one damaging *de novo* variant was determined, by subtracting the expected rate of damaging *de novo* variants per individual.

The formula is as follows:

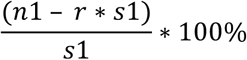

where n1 is the total number of sub-group CDH patients with at least one *de novo* deleterious variant, r is the expected rate per healthy individual with at least one *de novo* deleterious variant, where the rate was estimated by 10,000 simulations of Poisson distribution of variants per person, and s1 is the total number of sub-group CDH patients.

### Expression profile during diaphragm development

Mouse developing diaphragm (MDD) gene expression datasets from the pleuroperitoneal folds (PPFs)^33^ at embryonic day 11.5 (E11.5) were used in this study. High diaphragm expression is defined as the top quartile of probe sets based on RMA (Robust Multi-Array Average)-normalized expression levels of microarray data^19^.

### Single genes with multiple *de novo* mutations

For *MYRF*, the number of observed deleterious *de novo* mutations was compared to the expected deleterious mutation background using a Poisson test. The p-value passed Bonferroni correction with all protein-coding genes annotated in CCDS^34^.

## Results

### Clinical data of the cohort

Patients were recruited from the multicenter, longitudinal DHREAMS study ^35^ and from the Boston Children’s Hospital/Massachusetts General Hospital. In the combined cohort, there were 210 (59%) male and 147 (41%) female CDH patients. The gender distribution with increase male prevalence (1.4:1) is consistent with published retrospective and prospective studies ^**36**; **37**^. Among the 148 complex cases, the most frequent anomalies were congenital heart disease (41%), but neurodevelopmental delay, gastrointestinal, and other malformations were common (Table 1 and Supplementary Table 2). A total of 209 (59%) patients had isolated CDH without additional anomalies at last contact^20^. In the DHREAMS cohort (Online Methods) of 283 patients, 229 were part of the neonatal cohort (with 56% males), of which 152 had formal neurodevelopmental assessments at 2 years and/or 5 years. Nine (5.9%) patients evaluated had neurodevelopmental delay (NDD) with scores greater than 2 standard deviations below the mean (Supplementary Table 2).

**Table 1.** Clinical and phenotypic summary of CDH patients (n=357)

| Characteristics | Number | Percentage (%) |
| --- | --- | --- |
| **Male/Female** | 210/147 | 59/41 |
| **Left/Right/Other CDH location** | 269/56/32 | 75/16/9 |
| **White/Asian/Black/Other or unknown** | 240/13/10/94 | 67/4/3/26 |
| **Isolated cases** | 209 | 59 |
| **Complex cases** | 148 | 41 |
| congenital heart disease | 60 | 41 |
| gastrointestinal anomaly | 14 | 10 |
| structural brain anomaly | 15 | 10 |
| other congenital malformations | 67 | 45 |
| neurodevelopmental delay | 14 | 10 |

### Significant enrichment of coding *de novo* variants in both complex and isolated CDH

We identified 461 protein-coding *de novo* variants (Supplementary Table 3) (~1.29 per patient), including 190 damaging *de novo* variants in LGD and predicted deleterious missense variants (“D-mis” defined as CADD score ≥ 25, Supplementary Table 4). The overall *de novo* frequency in cases was 1.33 (255/192) in WGS and 1.25 (206/165) in WES. 41.2% (147/357) of probands carried at least one damaging *de novo* variant, including one *de novo* LGD in 8.4% (30/357), one *de novo* D-mis in 22.7% (81/357), and two or more damaging *de novos* in 10.1% (36/357).

We observed an overall enrichment of damaging *de novo* variants (fold enrichment (FE)=1.7, P=4.2×10^−4^ for LGD, and FE=1.5, P=3.2×10^−6^ for D-mis, respectively) in all CDH patients based on the expected mutation rate calibrated by the method described in Samocha et al.^29; 32^(Table 2, Online Methods). The positive predictive value (PPV) estimated from the enrichment rate for LGD and D-mis variants is 35%, which indicates about 67 damaging *de novo* variants contribute to CDH. The enrichment is still significant when stratifying complex and isolated CDH or by sex (Table 2). 22% of complex and 16% of isolated cases are explained by damaging *de novo* variants.

**Table 2.**
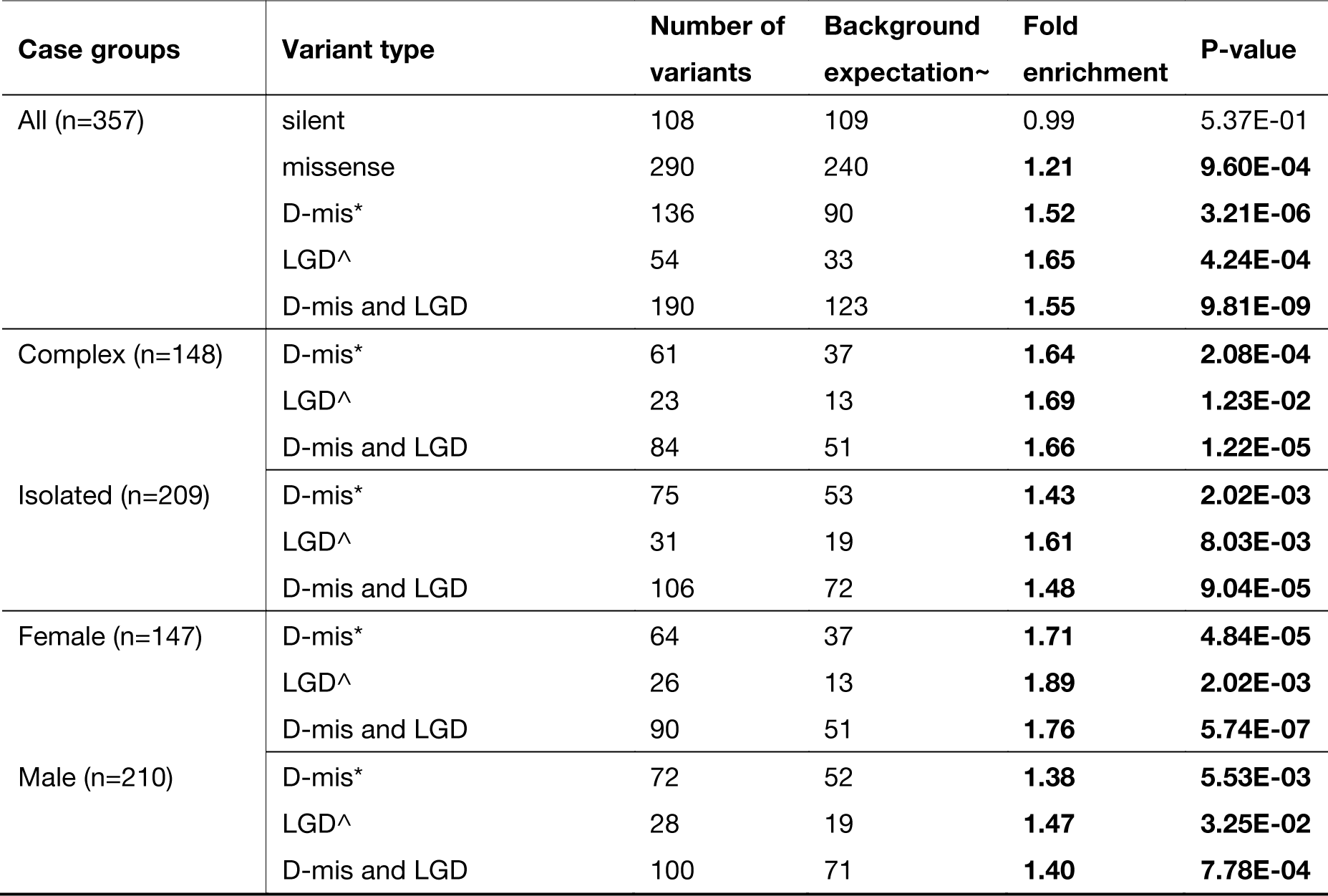
Enrichment of *de novo* variants in cases. ^LGD: likely-gene-disrupting, including frameshift, stopgain, stoploss, and splicing variants; *D-mis: missense predicted to be damaging by CADD phred score >= 25; ~Background expectation calibrated based on Samocha et al 2014 and Ware et al 2015^29; 32^.

We then tested whether the burden of damaging *de novo* variants were concentrated in constrained genes (defined as ExAC ^38^ pLI≥0.5)^38^ across variant types and sub-phenotypes. Overall, the burden of LGD variants was concentrated in constrained genes for both complex and isolated cases. The burden of D-mis variants was concentrated in constrained genes for complex cases, whereas for isolated cases, the burden of D-mis variants was concentrated in other genes (pLI<0.5 or not available) (Supplementary Table 5 and 6). This suggests that *de novo* pathogenic variants in constrained genes are more likely to cause syndromic abnormalities while such variants in other genes are more likely to cause isolated cases. Since other genes are generally not dosage sensitive, the observed burden of D-mis in these genes suggests a role of dominant negative or gain of function in isolated CDH.

### Different contribution of *de novo* variants to male and female CDH cases

Although CDH is more common in males, the enrichment of damaging *de novo* variants is higher in females than in males (FE=1.8 in female, FE=1.4 in male) (Table 2). We estimated that 27% of females can be explained by LGD or D-mis variants compared to 14% of males. In female cases, the enrichment rate of LGD or D-mis is comparable between complex and isolated cases (Supplementary Table 7). In contrast, in male cases, the enrichment rate is much higher in complex cases than isolated cases. In fact, there is essentially no enrichment of LGD or D-mis variants in male isolated cases (Figure 1a and Supplementary Table 7). Furthermore, in isolated female cases, LGD variants are mainly enriched in constrained genes (FE=3.3, P=0.001, Figure 1a), and D-mis variants were mainly in other genes (FE=2.2, P=0.0002) (Supplementary Table 8, Figure 1a). In complex CDH, the difference in enrichment rate of LGD and D-mis *de novo* variants in constrained genes between female and male cases is much smaller; and there is no significant enrichment of D-mis in other genes in either female or male cases (Supplementary Table 8, Figure 1b).

**Figure 1.**
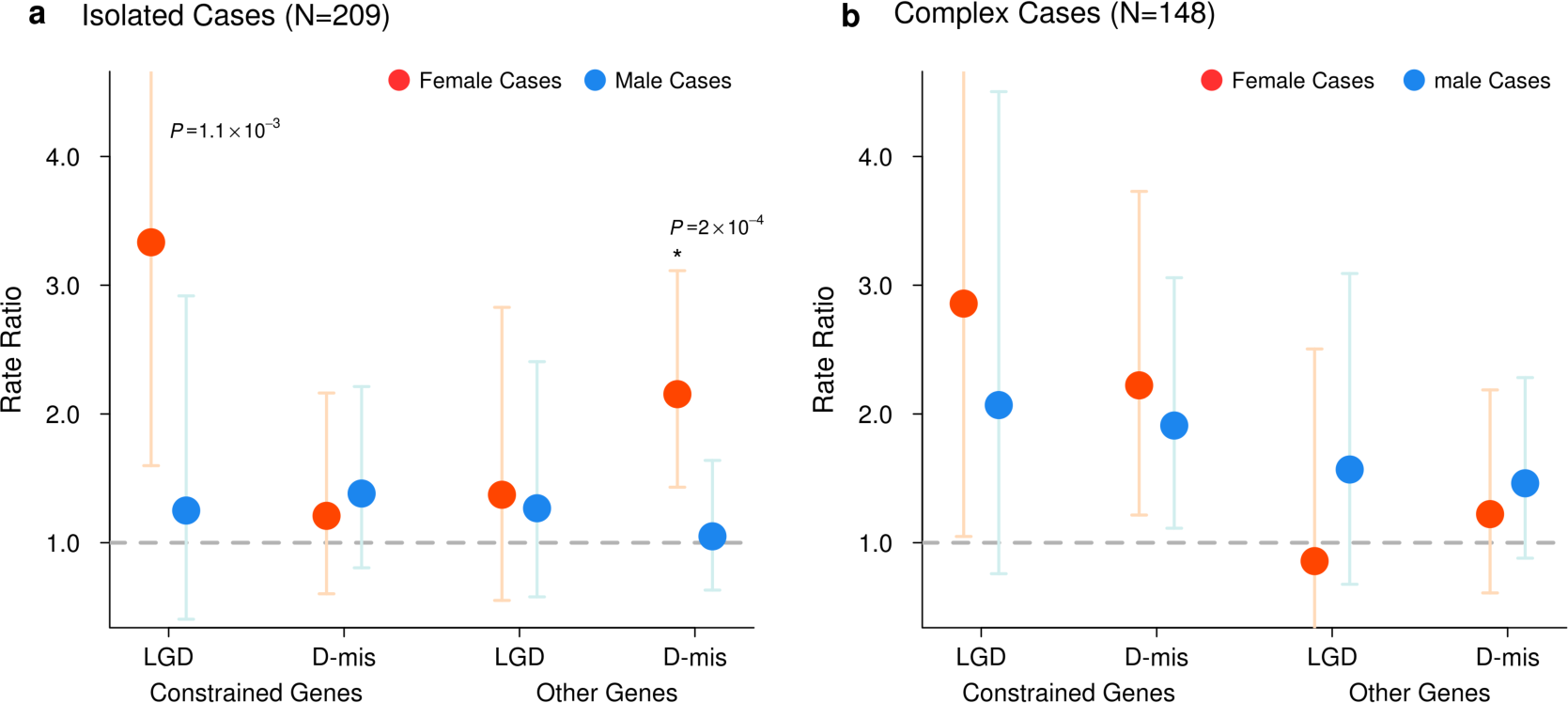
Female and male CDH cases have different enrichment rate of damaging de novo variants. (a) Enrichment of LGD variants and D-mis in constrained or other genes in isolated female and male cases. Constrained genes with LGD variants and other genes with D-mis variants are mainly enriched in female isolated cases. There is no enrichment of damaging de novo variants in isolated male cases. (b) Enrichment of LGD and D-mis variants in constrained or other genes in complex female and male cases. Both LGD and D-mis de novo variants were mainly enriched in constrained genes in complex cases. P-values shown are from tests of enrichment analysis. Red dots represent female cases, blue dots represent male cases. Bars represent the 95% confidence intervals (CIs) of the point estimates. Constrained genes: genes with ExAC pLI≥0.5. Other genes: genes with pLI<0.5 or no pLI estimate from ExAC; D-mis are missense variants with CADD Phred score≥25.

### Genes implicated by *de novo* LGD variants in complex and isolated CDH cases have distinct expression patterns in early diaphragm development

Genes associated with CDH are often expressed in pleuroperitoneal folds (PPF), an early structure critical in the developing diaphragm^39,10^. We analyzed the expression patterns of genes with LGD and D-mis variants using a mouse E11.5 PPF data set^33^. Isolated and complex cases have different patterns of LGD and missense variant burden. In complex cases, LGD *de novo* variants are dramatically enriched in genes in the top quartile of expression in developing diaphragm (E11.5) (FE=4.7, p=7×10^−7^) (Supplementary Table 9, Fig. 2). By contrast, in isolated cases, the burden of LGD *de novo* variants is distributed across genes with a broad range of expression in PPF (Supplementary Table 9 and 10, Fig. 2).

**Figure 2.**
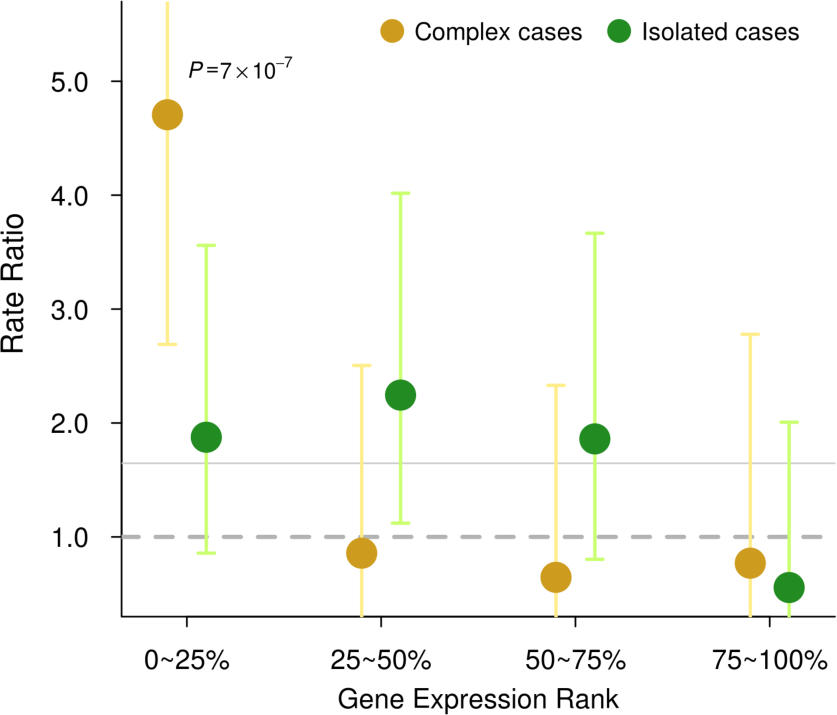
Isolated and complex cases have different enrichment patterns of LGD *de novo* variants. Enrichment rate of LGD *de novo* variants are shown in gene sets grouped by expression rank in E11.5 pleuroperitoneal folds (PPFs). In complex CDH cases, LGD *de novo* variants are dramatically enriched in the genes within the top quartile (0–25%) of expression in developing diaphragm (E11.5), and show no trend of enrichment in other quartiles. In isolated cases, LGD *de novo* variants have similar enrichment (~2x) across the 0–75% range of PPF gene expression. *P* values shown are from a test of enrichment. Bars represent the 95% CIs of the point estimates.

### *MYRF* is a novel candidate risk gene of CDH

Two genes are observed with multiple damaging *de novo* variants. Wilms tumor 1 (*WT1)* has been previously implicated in CDH^40^ and has two D-mis variants. Myelin Regulatory Factor (*MYRF)*, a transcription factor, has one *de novo* LGD and three D-mis variants (Fig. 3) in four complex CDH patients (p=2×10^−10^, based on comparison to expectation from background mutations ^29; 32^) (Table 3). A recent study of congenital heart disease (CHD) ^41; 42^ reported three additional damaging *de novo* missense variants (p.F387S, p.Q403H and p.L479V) in *MYRF* (Table 3, Fig 3a). All four CDH patients had CHD (Table 3). The CHD patient with the *MYRF* p.Q403H variant had hemidiaphragm eventration. Genitourinary anomalies were present in six of the seven patients, a female had a blind-ending vagina with no internal sex organs and five males had ambiguous genitalia or undescended testes. *MYRF* is a constrained gene intolerant of loss of function variants in the general populations (ExAC^38^ pLI=1). Although it has not previously been implicated in CDH or CHD, it is highly expressed in developing diaphragm and heart (ranked top 21% and 14% in mice E11.5 PPF ^33^ and E14.5 heart ^43^, respectively). Genital malformation may share developmental processes^44^ because PPF is physically connected dorsally to urogenital ridge.

**Table 3.** *De novo* variants of *MYRF* identified in CDH and CHD patients. Abbreviation: CDH (congenital diaphragmatic hernia); CHD (congenital heart disease); ASD (Atrial Septal Defect); VSD (Ventricular septal defect); TOF (Tetralogy of Fallot). The last three patients are ascertained by CHD as describe in Jin et al 2017 45

| Sample ID | Sex | Diaphragm defect | Heart defect | Genital defect | Other malformations | Protein | CADD |
| --- | --- | --- | --- | --- | --- | --- | --- |
| 01-1008 | Male | CDH | ASD,VSD,TOF | bilateral undescended testes | NA | p.G81Wfs*45 | 27.3 |
| 01-0429 | Female | CDH | VSD | no internal genital organs, external blind-ending vagina | accessory spleen | p.G435R | 32 |
| 04-0042 | Male | CDH | ASD,VSD | NA | NA | p.V679A | 25.9 |
| 05-0050 | Male | CDH | hypoplastic left heart syndrome | ambiguous genitalia | intellectual disability | p.R695H | 34 |
| 1-02264 | Male | NA | abnormal aorta | ambiguous genitalia, hypospadias, undescended testis | NA | p.F387S | 27.9 |
| 1-03160 | Male | right hemi-diaphragm eventration | abnormal atrial septum, pulmonary vein and arteries, systemic vein, aorta, aortic valve, mitral valve, ventricular septum | undescended testis | lung hypoplasia, abdominal abnormalities | p.Q403H | 27.6 |
| 1-07403 | Female | NA | abnormal aorta and aortic valve | NA | skeletal abnormalities, short stature | p.L479V | 23.9 |

**Figure 3.**
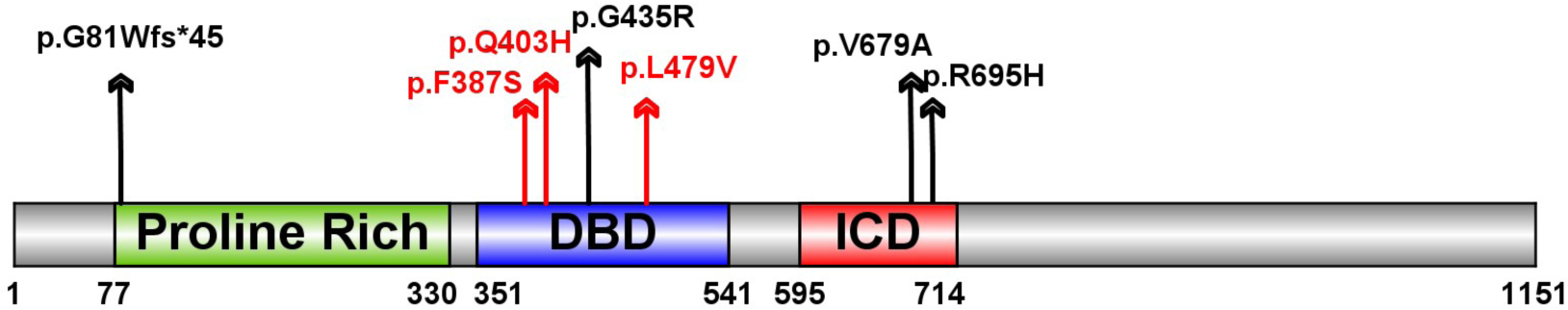
*De novo* variants identified in *MYRF*. Schematic of the MYRF protein with predicted sequence features, including N-terminal Proline Rich region, DNA-binding domain (DBD) and intramolecular chaperone domain (ICD). Variants identified in CDH indicated as black arrow, variants identified in congenital heart disease cases indicated with red arrows.

The three variants identified in CHD patients and p.G435R are located in the conserved DNA binding domain (DBD) of *MYRF* (Fig. 3), and could alter DNA binding^46^. The other two D-mis variants (p.V679R and p.R695H) are located in the intramolecular chaperone auto-processing domain (ICD) in a leucine zipper^47^. Mutations in the leucine zipper of the ICD domain may inhibit the trimerization of MYRF, resulting in the failure of formation of the N-terminal trimer^47^ which is important for the transcription factor function^48^. MYRF is thought to be an essential transcription factor for oligodendrocyte differentiation and myelination^49^. Conditional deletion of *Myrf* impaired motor learning^50; 51^ and the individual with the p.V679A variant we assessed at two years old had intellectual disability.

## Discussions

CDH is slightly more common in males. For the first time, our study suggests that male and female CDH may have a different genetic architecture, especially among isolated CDH cases. Damaging *de novo* variants with large effect have a substantial contribution to isolated female cases but little contribution to isolated male cases. Given the higher frequency of males among isolated cases, a plausible explanation is that polygenic risk from inherited variants alone can cause isolated CDH in males, but due to a female protective effect^52^, additional highly penetrant *de novo* variants are often required to cause CDH in females to pass the threshold of liability. This is similar to what has been observed in autism which is also more common among males ^53^. Since there is a similar male/female ratio in overall cohort and our neonatal cohort (1.4:1), this difference is unlikely due to ascertainment or survival bias. The parental ages for male and female probands were similar and cannot account for the differences we observed in *de novo* variants.

Additionally, we found genes implicated in isolated and complex cases have distinct expression patterns in early development. In complex CDH, the burden of LGD and D-mis variants are concentrated in genes highly expressed in the FFP, an early embryonic diaphragm precursor, consistent with the pleiotropic effects of these genes on diaphragm and other organogenesis. By contrast, the burden of LGD variants in isolated cases is distributed across genes with a broader range of expression in PPF. Since the bulk expression data from PPFs is the sum of different cell types^54^, the lack of correlation of LGD enrichment and expression level in PPF suggests the possibility that a substantial portion of the implicated genes in isolated cases could be expressed only in sub-populations of cells in the PPF that are not relevant to organogenesis in other parts of the body. Single-cell mRNA-sequencing will be necessary to analyze gene expression pattern in specific cell types and further assess the cell type(s) responsible for isolated CDH.

Finally, *MYRF* is a novel candidate risk gene of CDH. The four CDH patients carrying damaging *de novo* variants in *MYRF* all have congenital heart defects, genitourinary anomalies including ambiguous genital, and this likely represents a novel syndrome. As we identify larger numbers of patients with mutations in genes associated with CDH, we will be able to better describe the spectrum of disease associated with the gene as well as the clinical outcomes including risk of pulmonary hypertension and respiratory complications which are life threatening concerns for CDH patients. Identification of additional high risk CDH genes should elucidate the developmental biology and provide targets for treatment and prevention.

## Acknowledgements

We would like to thank the patients and their families for their generous contribution. We are grateful for the technical assistance provided by Patricia Lanzano, Jiancheng Guo, and Liyong Deng from Columbia University, Jessica Kim at Boston Children’s Hospital, and Caroline Coletti and Pooja Bhayani at Massachusetts General Hospital. We thank our clinical coordinators across the DHREAMS centers: Trish Burns at Cincinnati Children's Hospital, Sheila Horak at Children's Hospital & Medical Center of Omaha, Brandy Gonzales at Oregon Health and Science University, Karen Lukas at St. Louis Children's Hospital, Jeannie Kreutzman at CS Mott Children's Hospital, Min Shi at Children's Hospital of Pittsburgh, Michelle Knezevich and Cheryl Kornberg at Medical College of Wisconsin. We thank University of Washington Center for Mendelian Genomics (UW-CMG) team for generating part of the WES data. The WES data generation at UW-CMG was funded by the National Human Genome Research Institute and the National Heart, Lung and Blood Institute grant HG006493 to Drs. Debbie Nickerson, Michael Bamshad, and Suzanne Leal. The whole genome sequencing data were generated by NIH Gabriella Miller Kids First Pediatric Research Program (X01HL132366). This work was supported by NIH grants R01HD057036 (L.Y., J.W., W.K.C.), R03HL138352 (A.K., W.K.C., Y.S.), R01GM120609 (H.Q., Y.S.), UL1 RR024156 (W.K.C.), and 1P01HD068250 (P.K.D, M.L., F.A.H., J.M.W., W.K.C.) Additional funding support was provided by grant from CHERUBS, a grant from the National Greek Orthodox Ladies Philoptochos Society, Inc. and generous donations from The Wheeler foundation, Vanech Family Foundation, Larsen Family, Wilke Family and many other families.

## Author Contributions

W.K.C and Y.S. conceived the study. F.A.H., J.M.W., and P.K.D. provided genomic data. H.Q., X.Z., Y.L., A.K., W.K.C., and Y.S. analyzed and interpreted the data. Y.L., X.Z., H.Q., W.K.C., and Y.S. wrote the manuscript. J.W., G.A., F.L., T.C., R.C., K.A., M.E.D., D.C., B.W.W., G.B.M., D.P., A.J.W., M.E., F.A.H., M.L., J.M.W., P.K.D. collected samples and clinical information. All authors contributed and discussed the results and critically reviewed the manuscript.

## References

1. Chandrasekharan, P.K., Rawat, M., Madappa, R., Rothstein, D.H., and Lakshminrusimha, S. (2017). Congenital Diaphragmatic hernia - a review. Matern Health Neonatol Perinatol 3, 6.

2. Pober, B.R., Russell, M.K., and Ackerman, K.G. (1993). Congenital Diaphragmatic Hernia Overview. In GeneReviews(R), R.A. Pagon, M.P. Adam, H.H. Ardinger, S.E. Wallace, A. Amemiya, L.J.H. Bean, T.D. Bird, N. Ledbetter, H.C. Mefford, R.J.H. Smith, et al., eds. (Seattle (WA)).

3. Pober, B.R. (2007). Overview of epidemiology, genetics, birth defects, and chromosome abnormalities associated with CDH. Am J Med Genet C Semin Med Genet 145C, 158–171.

4. Danzer, E., Gerdes, M., Bernbaum, J., D'Agostino, J., Bebbington, M.W., Siegle, J., Hoffman, C., Rintoul, N.E., Flake, A.W., Adzick, N.S., et al. (2010). Neurodevelopmental outcome of infants with congenital diaphragmatic hernia prospectively enrolled in an interdisciplinary follow-up program. J Pediatr Surg 45, 1759–1766.

5. Graziano, J.N., and Congenital Diaphragmatic Hernia Study, G. (2005). Cardiac anomalies in patients with congenital diaphragmatic hernia and their prognosis: a report from the Congenital Diaphragmatic Hernia Study Group. J Pediatr Surg 40, 1045–1049; discussion 1049–1050.

6. Skari, H., Bjornland, K., Haugen, G., Egeland, T., and Emblem, R. (2000). Congenital diaphragmatic hernia: a meta-analysis of mortality factors. J Pediatr Surg 35, 1187–1197.

7. Zalla, J.M., Stoddard, G.J., and Yoder, B.A. (2015). Improved mortality rate for congenital diaphragmatic hernia in the modern era of management: 15 year experience in a single institution. J Pediatr Surg 50, 524–527.

8. Leeuwen, L., and Fitzgerald, D.A. (2014). Congenital diaphragmatic hernia. J Paediatr Child Health 50, 667–673.

9. Ackerman, K.G., and Greer, J.J. (2007). Development of the diaphragm and genetic mouse models of diaphragmatic defects. Am J Med Genet C Semin Med Genet 145C, 109–116.

10. Merrell, A.J., Ellis, B.J., Fox, Z.D., Lawson, J.A., Weiss, J.A., and Kardon, G. (2015). Muscle connective tissue controls development of the diaphragm and is a source of congenital diaphragmatic hernias. Nat Genet 47, 496–504.

11. Jay, P.Y., Bielinska, M., Erlich, J.M., Mannisto, S., Pu, W.T., Heikinheimo, M., and Wilson, D.B. (2007). Impaired mesenchymal cell function in Gata4 mutant mice leads to diaphragmatic hernias and primary lung defects. Dev Biol 301, 602–614.

12. Ackerman, K.G., Herron, B.J., Vargas, S.O., Huang, H.L., Tevosian, S.G., Kochilas, L., Rao, C., Pober, B.R., Babiuk, R.P., Epstein, J.A., et al. (2005). Fog2 is required for normal diaphragm and lung development in mice and humans. Plos Genet 1, 58–65.

13. Castiglia, L., Fichera, M., Romano, C., Galesi, O., Grillo, L., Sturnio, M., and Failla, P. (2005). Narrowing the candidate region for congenital diaphragmatic hernia in chromosome 15q26: Contradictory results. Am J Hum Genet 77, 892–894.

14. Klaassens, M., van Dooren, M., Eussen, H.J., Douben, H., den Dekker, A.T., Lee, C., Donahoe, P.K., Galjaard, R.J., Goemaere, N., de Krijger, R.R., et al. (2005). Congenital Diaphragmatic hernia and chromosome 15q26: Determination of a candidate region by use of fluorescent in situ hybridization and array-based comparative genomic hybridization. Am J Hum Genet 76, 877–882.

15. Shimokawa, O., Miyake, N., Yoshimura, T., Sosonkina, N., Harada, N., Mizuguchi, T., Kondoh, S., Kishino, T., Ohta, T., Remco, V., et al. (2005). Molecular characterization of del(8)(p23.1p23.1) in a case of congenital diaphragmatic hernia. American Journal of Medical Genetics Part A 136A, 49–51.

16. Wat, M.J., Shchelochkov, O.A., Holder, A.M., Breman, A.M., Dagli, A., Bacino, C., Scaglia, F., Zori, R.T., Cheung, S.W., Scott, D.A., et al. (2009). Chromosome 8p23.1 Deletions as a Cause of Complex Congenital Heart Defects and Diaphragmatic Hernia. American Journal of Medical Genetics Part A 149A, 1661–1677.

17. You, L.R., Takamoto, N., Yu, C.T., Tanaka, T., Kodama, T., DeMayo, F.J., Tsai, S.Y., and Tsai, M.J. (2005). Mouse lacking COUP-TFII as an animal model of Bochdalek-type congenital diaphragmatic hernia. Proceedings of the National Academy of Sciences of the United States of America 102, 16351–16356.

18. Longoni, M., High, F.A., Qi, H., Joy, M.P., Hila, R., Coletti, C.M., Wynn, J., Loscertales, M., Shan, L., Bult, C.J., et al. (2017). Genome-wide enrichment of damaging de novo variants in patients with isolated and complex congenital diaphragmatic hernia. Hum Genet 136, 679–691.

19. Yu, L., Sawle, A.D., Wynn, J., Aspelund, G., Stolar, C.J., Arkovitz, M.S., Potoka, D., Azarow, K.S., Mychaliska, G.B., Shen, Y., et al. (2015). Increased burden of de novo predicted deleterious variants in complex congenital diaphragmatic hernia. Hum Mol Genet 24, 4764–4773.

20. Yu, L., Wynn, J., Ma, L., Guha, S., Mychaliska, G.B., Crombleholme, T.M., Azarow, K.S., Lim, F.Y., Chung, D.H., Potoka, D., et al. (2012). De novo copy number variants are associated with congenital diaphragmatic hernia. J Med Genet 49, 650–659.

21. Li, H., and Durbin, R. (2009). Fast and accurate short read alignment with Burrows-Wheeler transform. Bioinformatics 25, 1754–1760.

22. Li, H., Ruan, J., and Durbin, R. (2008). Mapping short DNA sequencing reads and calling variants using mapping quality scores. Genome research 18, 1851–1858.

23. GATK-Team. (2016). GATK best practice.

24. Price, A.L., Patterson, N.J., Plenge, R.M., Weinblatt, M.E., Shadick, N.A., and Reich, D. (2006). Principal components analysis corrects for stratification in genome-wide association studies. Nature genetics 38, 904–909.

25. International HapMap, C., Altshuler, D.M., Gibbs, R.A., Peltonen, L., Altshuler, D.M., Gibbs, R.A., Peltonen, L., Dermitzakis, E., Schaffner, S.F., Yu, F., et al. (2010). Integrating common and rare genetic variation in diverse human populations. Nature 467, 52–58.

26. Purcell, S., Neale, B., Todd-Brown, K., Thomas, L., Ferreira, M.A., Bender, D., Maller, J., Sklar, P., de Bakker, P.I., Daly, M.J., et al. (2007). PLINK: a tool set for whole-genome association and population-based linkage analyses. American journal of human genetics 81, 559–575.

27. Chang, C.C., Chow, C.C., Tellier, L.C., Vattikuti, S., Purcell, S.M., and Lee, J.J. (2015). Second-generation PLINK: rising to the challenge of larger and richer datasets. GigaScience 4, 7.

28. Homsy, J., Zaidi, S., Shen, Y., Ware, J.S., Samocha, K.E., Karczewski, K.J., DePalma, S.R., McKean, D., Wakimoto, H., Gorham, J., et al. (2015). De novo mutations in congenital heart disease with neurodevelopmental and other congenital anomalies. Science 350, 1262–1266.

29. Samocha, K.E., Robinson, E.B., Sanders, S.J., Stevens, C., Sabo, A., McGrath, L.M., Kosmicki, J.A., Rehnstrom, K., Mallick, S., Kirby, A., et al. (2014). A framework for the interpretation of de novo mutation in human disease. Nat Genet 46, 944–950.

30. Wang, K., Li, M., and Hakonarson, H. (2010). ANNOVAR: functional annotation of genetic variants from high-throughput sequencing data. Nucleic acids research 38, e164.

31. Kircher, M., Witten, D.M., Jain, P., O'Roak, B.J., Cooper, G.M., and Shendure, J. (2014). A general framework for estimating the relative pathogenicity of human genetic variants. Nat Genet 46, 310–315.

32. Ware, J.S., Samocha, K.E., Homsy, J., and Daly, M.J. (2015). Interpreting de novo Variation in Human Disease Using denovolyzeR. Current protocols in human genetics / editorial board, Jonathan L Haines [et al] 87, 7 25 21–27 25 15.

33. Russell, M.K., Longoni, M., Wells, J., Maalouf, F.I., Tracy, A.A., Loscertales, M., Ackerman, K.G., Pober, B.R., Lage, K., Bult, C.J., et al. (2012). Congenital diaphragmatic hernia candidate genes derived from embryonic transcriptomes. Proc Natl Acad Sci U S A 109, 2978–2983.

34. Pruitt, K.D., Harrow, J., Harte, R.A., Wallin, C., Diekhans, M., Maglott, D.R., Searle, S., Farrell, C.M., Loveland, J.E., Ruef, B.J., et al. (2009). The consensus coding sequence (CCDS) project: Identifying a common protein-coding gene set for the human and mouse genomes. Genome Res 19, 1316–1323.

35. DHREAMS: Diaphragmatic Hernia Research & Exploration; Advancing Molecular Science, http://www.cdhgenetics.com/. (2009).

36. Hinton, C.F., Siffel, C., Correa, A., and Shapira, S.K. (2017). Survival Disparities Associated with Congenital Diaphragmatic Hernia. Birth Defects Res 109, 816–823.

37. Leeuwen, L., Mous, D.S., van Rosmalen, J., Olieman, J.F., Andriessen, L., Gischler, S.J., Joosten, K.F.M., Wijnen, R.M.H., Tibboel, D., H, I.J., et al. (2017). Congenital Diaphragmatic Hernia and Growth to 12 Years. Pediatrics 140.

38. Lek, M., Karczewski, K.J., Minikel, E.V., Samocha, K.E., Banks, E., Fennell, T., O'Donnell-Luria, A.H., Ware, J.S., Hill, A.J., Cummings, B.B., et al. (2016). Analysis of protein-coding genetic variation in 60,706 humans. Nature 536, 285–291.

39. Clugston, R.D., Zhang, W., and Greer, J.J. (2008). Gene expression in the developing diaphragm: significance for congenital diaphragmatic hernia. Am J Physiol Lung Cell Mol Physiol 294, L665–675.

40. Carmona, R., Canete, A., Cano, E., Ariza, L., Rojas, A., and Munoz-Chapuli, R. (2016). Conditional deletion of WT1 in the septum transversum mesenchyme causes congenital diaphragmatic hernia in mice. Elife 5.

41. Homsy, J., Zaidi, S., Shen, Y., Ware, J.S., Samocha, K.E., Karczewski, K.J., DePalma, S.R., McKean, D., Wakimoto, H., Gorham, J., et al. (2015). De novo mutations in congenital heart disease with neurodevelopmental and other congenital anomalies. Science (New York, NY) 350, 1262–1266.

42. Jin, S.C., Homsy, J., Zaidi, S., Qi, H., Lu, Q., Zeng, X., Chung, W.-C., Shen, Y., Zhao, H., Kaltman, J.R., et al. (2017). Contribution of rare inherited and de novo variants in 2,871 congenital heart disease probands. Nature Genetics In press.

43. Zaidi, S., Choi, M., Wakimoto, H., Ma, L., Jiang, J., Overton, J.D., Romano-Adesman, A., Bjornson, R.D., Breitbart, R.E., Brown, K.K., et al. (2013). De novo mutations in histone-modifying genes in congenital heart disease. Nature 498, 220–223.

44. Azarow, K.S., Cusick, R., Wynn, J., Chung, W., Mychaliska, G.B., Crombleholme, T.M., Chung, D.H., Lim, F.Y., Potoka, D., Warner, B.W., et al. (2015). The association between congenital diaphragmatic hernia and undescended testes. J Pediatr Surg 50, 744–745.

45. Jin, S.C., Homsy, J., Zaidi, S., Lu, Q., Morton, S., DePalma, S.R., Zeng, X., Qi, H., Chang, W., Sierant, M.C., et al. (2017). Contribution of rare inherited and de novo variants in 2,871 congenital heart disease probands. Nat Genet.

46. Bujalka, H., Koenning, M., Jackson, S., Perreau, V.M., Pope, B., Hay, C.M., Mitew, S., Hill, A.F., Lu, Q.R., Wegner, M., et al. (2013). MYRF is a membrane-associated transcription factor that autoproteolytically cleaves to directly activate myelin genes. PLoS Biol 11, e1001625.

47. Li, Z., Park, Y., and Marcotte, E.M. (2013). A Bacteriophage tailspike domain promotes self-cleavage of a human membrane-bound transcription factor, the myelin regulatory factor MYRF. PLoS Biol 11, e1001624.

48. Kim, D., Choi, J.O., Fan, C., Shearer, R.S., Sharif, M., Busch, P., and Park, Y. (2017). Homo-trimerization is essential for the transcription factor function of Myrf for oligodendrocyte differentiation. Nucleic Acids Res 45, 5112–5125.

49. Hornig, J., Frob, F., Vogl, M.R., Hermans-Borgmeyer, I., Tamm, E.R., and Wegner, M. (2013). The transcription factors Sox10 and Myrf define an essential regulatory network module in differentiating oligodendrocytes. PLoS Genet 9, e1003907.

50. McKenzie, I.A., Ohayon, D., Li, H., de Faria, J.P., Emery, B., Tohyama, K., and Richardson, W.D. (2014). Motor skill learning requires active central myelination. Science 346, 318–322.

51. Xiao, L., Ohayon, D., McKenzie, I.A., Sinclair-Wilson, A., Wright, J.L., Fudge, A.D., Emery, B., Li, H., and Richardson, W.D. (2016). Rapid production of new oligodendrocytes is required in the earliest stages of motor-skill learning. Nat Neurosci 19, 1210–1217.

52. Robinson, E.B., Lichtenstein, P., Anckarsater, H., Happe, F., and Ronald, A. (2013). Examining and interpreting the female protective effect against autistic behavior. Proc Natl Acad Sci U S A 110, 5258–5262.

53. Iossifov, I., O'Roak, B.J., Sanders, S.J., Ronemus, M., Krumm, N., Levy, D., Stessman, H.A., Witherspoon, K.T., Vives, L., Patterson, K.E., et al. (2014). The contribution of de novo coding mutations to autism spectrum disorder. Nature 515, 216–221.

54. Clugston, R.D., and Greer, J.J. (2007). Diaphragm development and congenital diaphragmatic hernia. Semin Pediatr Surg 16, 94–100.

